# Adaptations and responses of the common dandelion to low atmospheric pressure in high altitude environments

**DOI:** 10.1101/2021.01.28.428657

**Authors:** Carla C. M. Arce, Zoe Bont, Ricardo A. R. Machado, Paulo F. Cristaldo, Matthias Erb

**Author notes:** **Correspondence:** Matthias Erb, University of Bern, Institute of Plant Sciences, Altenbergrain 21, 3013 Bern, Switzerland. shared first authorship (Carla C. M. Arce, Zoe Bont). **Statement of authorship:** ME, ZB and CA designed the study. CA, RM and ZB collected data. ZB, CA and PF analysed and interpreted the data. ZB and ME wrote the first draft of the manuscript. All authors contributed to the final version of the manuscript. **Data accessibility statement:** All data supporting this study will be stored in the Dryad Digital Repository and the data DOI will be included in the manuscript.

## Abstract

Atmospheric pressure is an important, yet understudied factor that may shape plant ecology and evolution. By growing plants under controlled conditions at different experimental stations in the Swiss alps, we evaluated the impact of ecologically realistic atmospheric pressures between 660 and 950 hPa on the growth and defence of different dandelion populations. Low atmospheric pressure was associated with reduced root growth and defensive sesquiterpene lactone production. Defense suppression only occurred in populations originating from lower altitudes. Populations from higher altitudes constitutively produced less sesquiterpene lactones and did not suffer from suppression under low atmospheric pressure. We conclude that atmospheric pressure modulates root growth and defence traits, and that evolutionary history shapes plant phenotypic responses to atmospheric pressure. Our findings have important implications for our understanding of altitudinal gradients and the future use of plants as a source of food and bioactive metabolites in extraterrestrial habitats.

## Introduction

Plants successfully colonize a wide range of terrestrial habitats. Their capacity to survive under variable and challenging environmental conditions is a prerequisite for their contribution to biodiversity and sustainable agriculture in the face of climate change, and their potential use as a source of food and medicine in future extraterrestrial habitats (Jump & Peñuelas 2005; Wheeler 2017). Atmospheric pressure plays a potentially important role in terrestrial and extraterrestrial environments, albeit for different reasons. On earth, climate change research often relies on altitudinal gradients to understand how plants and terrestrial ecosystems will be affected by future climates. Such studies typically assume that differences in climate at different altitudes account for changes in plant performance, and that effects of atmospheric pressure are negligible. In contrast, space plant biology has long been interested in understanding effects of atmospheric pressure on plants, as it would be more practical to maintain low pressure greenhouses in future moon or mars colonies (Corey *et al.* 1997; Iwabuchi & Kurata 2003; Richards *et al.* 2006). Thus, understanding the impact of atmospheric pressure on plants and the capacity of plants to withstand and adapt to different atmospheric pressures is of substantial interest for the present and future of humanity. Yet, to date, our understanding of how atmospheric pressure influences plants is not well developed.

How are plants affected by low atmospheric pressure? One consequence of low atmospheric pressure at high altitudes is the decrease of partial pressures of O2 and CO2, which are essential substrates for respiration and photosynthesis (Zabalza *et al.* 2009; Xu *et al.* 2015). Changes in O_2_ and CO_2_ partial pressures have been linked to changes in plant physiology and growth (Paul *et al.* 2004; He *et al.* 2007; Kammer *et al.* 2015; Zhou *et al.* 2017). Effects are complex, since the decrease of the partial pressure of the atmospheric gases is accompanied by an increase in diffusion rates, which may compensate for the low ambient concentration of the essential gases (Terashima *et al.* 1995). Further, as the diffusion coefficient for water vapor is increased, transpiration increases, (Smith & Geller 1979), which can impose water stress on plants growing under reduced atmospheric pressure (Iwabuchi & Kurata 2003; Paul *et al.* 2004; Richards *et al.* 2006).

Can plants exhibit phenotypic plasticity to cope with low atmospheric pressure? Several studies suggest that plants respond dynamically to reduced atmospheric pressure (Spanarkel & Drew 2002; He *et al.* 2003; Iwabuchi & Kurata 2003; Richards *et al.* 2006). In the mountain plant *Arabis alpina*, for instance, stomatal density increases and stomata aperture narrows at low atmospheric pressure (Kammer *et al.* 2015). This induced response likely benefits the plant, because at low partial pressures of atmospheric gases, a higher stomatal density may ensure optimal supply of CO_2_ for photosynthesis (Woodward & Bazzaz 1988; Xu *et al.* 2016), while a narrow aperture of the stomata can restrict water loss to counteract increased transpiration rate (Buckley 2005). At a molecular level, it is assumed that low atmospheric pressure represents an environmental stress to which plants must respond with changes in their metabolic pathways in order to survive successfully (Ferl *et al.* 2002; Paul & Ferl 2006). Recent research has documented extensive changes in gene expression patterns in *Arabidopsis thaliana* when exposed to a low atmospheric pressure environment, including genes associated with hypoxia and water loss (Paul *et al.* 2004; Zhou *et al.* 2017).

Can plants adapt to low atmospheric pressure over evolutionary time? Vascular plants colonize habitats between 0 and 6150 m above sea corresponding to atmospheric pressures between 101 and 46 kPa (Angel *et al.* 2016). Since atmospheric pressure influences plant performance, and many plant species occur along altitudinal gradients, local adaptation to atmospheric pressure can be expected (Kammer *et al.* 2015; Ward & Strain 1997; Ward *et al.* 2000). Using growth chambers simulating high and low altitude pressure conditions, Kammer *et al*. (2015) found that *Arabis alpina* adjusts stomatal density in response to low atmospheric pressure, while the low altitude plant *Arabidopsis thaliana* does not. Evidence for genetic differences in the response of stomatal density to pressure conditions was also found within species. Woodward & Bazzaz (1988) showed that in the grass *Nardus stricta*, plants from higher altitudes developed greater declines in stomatal density at experimentally increased CO_2_ partial pressure than plants from lower altitudes. Apart from stomatal development, very little is known about evolutionary adaptations of plants to low atmospheric pressure.

Plant secondary or specialized metabolites play important roles in plant responses and adaptations to diverse environments and stress factors (Hartmann 2007; Moore *et al.* 2014), including herbivores and pathogens (Ehrlich & Raven 1964; Kessler & Baldwin 2001; Bednarek & Osbourn 2009; Moles *et al.* 2013), abiotic stress (Ramakrishna & Ravishankar 2011; Arbona *et al.* 2013; Nakabayashi & Saito 2015), mutualists (Pichersky & Gershenzon 2002; Schliemann *et al.* 2008; Stevenson *et al.* 2017) and other plants (Baldwin *et al.* 2006; Semchenko *et al.* 2014). The production, transport and storage of specialized metabolites is assumed to be costly (Neilson *et al.* 2013), and plants therefore constantly fine-tune their chemical arsenal to the demands of their environment and internal condition through phenotypic plasticity (Metlen *et al.* 2009). Over evolutionary times, environmental conditions may act as selective forces on plant genotype selection and can shape genetically determined chemical profiles of plants (Cunningham *et al.* 1999; Agrawal *et al.* 2012; Züst *et al.* 2012; Kessler & Kalske 2018). With increasing altitude, mountain habitats impose different environmental demands on plants, including harsher abiotic conditions and a lower intensity of biotic interactions (Rasmann *et al.* 2014; Buckley *et al.* 2019; Midolo & Wellstein 2020). Several studies found evidence that the genetic variation of plant secondary metabolites is shaped by these environmental gradients (Bernal *et al.* 2013; Moreira *et al.* 2018; Bakhtiari *et al.* 2019; Buckley *et al.* 2019; Bont *et al.* 2020). However, the role of decreasing atmospheric pressure at increasing altitudes in shaping the evolution and expression of plant secondary metabolites is poorly understood. Levine *et al.* (2008) found that the glucosinolate content in the roots of radish was decreased under hypobaric, normoxic conditions. In lettuce, the concentration of phenolics, anthocyanins and carotenoids in the leaves was increased under hypobaric and hypoxic conditions (He *et al.* 2013). However, all these experiments were conducted in low pressure chambers under conditions far beyond the natural range of plants. How a plant’s evolutionary history shapes its response to ecologically realistic low-pressure environments is unknown.

Along environmental gradients, closely related asexual and sexual plant taxa often have different distribution patterns – a phenomenon called geographic parthenogenesis (Glesener & Tilman 1978). It is generally predicted that apomictic plants with asexual reproduction tend to have larger distribution ranges including higher latitudes and higher altitudes, and thus lower atmospheric pressure, than their sexual relatives (Bierzychudek 1985; Kearney 2005; Cosendai *et al.* 2013). Hypotheses to explain these environmental distribution patterns include different colonization abilities of sexual and asexual organisms as well as different capacities to co-evolve with other organisms (e.g. reviewed in Hörandl 2006 and in Tilquin & Kokko 2016). Although many studies empirically support the predicted geographical parthenogenesis patterns, some taxa with asexual and sexual organisms show opposite or mixed trends in geographic distribution. For the common dandelion *Taraxacum officinale* agg. (Asteraceae), a species complex that largely consists of diploid sexuals and triploid asexuals (Verduijn *et al.* 2004), larger latitudinal distribution of triploids has been observed, with triploids colonizing more extreme environments in the north of Europe (Menken *et al.* 1995; Van Dijk *et al.* 2003; Verhoeven & Biere 2013). Surprisingly however, triploids are less frequent at higher altitudes than diploids, at least along certain transects (Calame & Felber 2000; Bont *et al.* 2020). The reason for this pattern is currently unclear. One possibility is that high altitude environments impose selection pressures that are different from high latitude environments. Therefore, one hypothesis to explain the lower success of triploids at high altitude is that they are constrained by low resistance to low atmospheric pressure.

*Taraxacum officinale* possesses a reservoir of secondary metabolites stored in specialized cells, so called laticifers, throughout almost all organs. The latex of *T. officinale* is most abundant in the taproot. The latex is characterized by three major classes of secondary metabolites: hydroxyphenylacetate inositol esters with either two or three side chains (di-PIEs and tri-PIEs), the sesquiterpene lactone taraxinic acid β-d-glucopyranosyl ester (TA-G), and triterpene acetates (TritAc) (Huber *et al.* 2015). Latex is mainly connotated with defensive functions against herbivores and pathogens (Konno 2011), and our previous work on the bioactivity and ecological role of the latex metabolites of *T. officinale* confirms this hypothesis (Huber *et al.* 2016b, a; Bont *et al.* 2017). TA-G in particular reduces the attractiveness of *T. officinale* to white grubs and thereby increases plant performance (Huber *et al.* 2016b). Interestingly, in our recent study on the latex metabolites of 63 natural *T. officinale* populations across Switzerland we found a strong association of the latex metabolites TA-G and di-PIEs with the climatic history of the natural populations, which may be suggestive of a role of latex secondary metabolites in abiotic stress tolerance (Bont *et al.* 2020). Inside the lacticifers, latex is maintained at positive pressure, which allows it to be expelled after tissue disruption. Latex likely responds to turgor pressure (Agrawal & Konno 2009), which in turn is determined by the water balance of the plant. At low atmospheric pressure, evaporation is increased, which could therefore have a direct or indirect effect on latex quality or quantity.

Here, we took advantage of experimental stations in Switzerland between 526 and 3450 m with standardized abiotic conditions to study the effects of atmospheric pressure on the growth and latex composition of *T. officinale*. We included offspring from nine natural populations from Switzerland from different altitudes, including populations containing both diploid and triploid cytotypes. This setup allowed us to test (i) whether low atmospheric pressure affects plant growth and latex composition through environmental plasticity; (ii) whether there are signatures of local adaptation to atmospheric pressure, with populations from higher altitudes performing better at low atmospheric pressure; and (iii) whether low atmospheric pressure has a stronger impact on triploid than diploid plants, thus potentially restricting the expansion of triploids towards higher altitudes.

## Material and methods

### Study species and seed collection

The common dandelion is a latex-producing species complex with a worldwide, cosmopolitan distribution (Stewart-Wade *et al.* 2002). The wind-dispersed perennial herb can tolerate a broad range of environmental conditions and can be found from sea level to altitudes of up to 4000 m.a.s.l. (Molina-Montenegro *et al.* 2013; Sandoya *et al.* 2017). In Switzerland, the plant occurs at altitudes of up to 2000 m.a.s.l. (Calame & Felber 2000). In this study, plants from nine natural populations of Switzerland were included. The nine populations are a subset of 63 populations that were characterized in our previous work (Bont *et al.* 2020). They cover an altitudinal range from 302 to 1607 m.a.s.l (Fig. 1ab, Table S1 in Supporting Information). Each population was located within a maximal distance of 1 km from a meteorological monitoring station of MeteoSwiss, the Swiss Federal Office for Meteorology and Climatology, which enabled us to obtain long-term data on climatic conditions. According to Bont *et al.* (2020), 10 variables were selected from the MeteoSwiss database, representing average air pressure, temperature, precipitation and light conditions of the populations for the years 1996 – 2015. For most variables, data was available for all 9 populations (Fig. S1c). Seeds from the natural populations were collected and F2 plants were generated for each population as described in Bont *et al.* 2020. The cytotype distribution of each population was determined by analysing the ploidy level of the plants with flow cytometry (Bont *et al.* 2020). The nine F2 populations consisted of populations with only diploid plants (5 populations) and populations with both triploid and diploid plants (4 populations) (Table S1).

**Figure 1:**
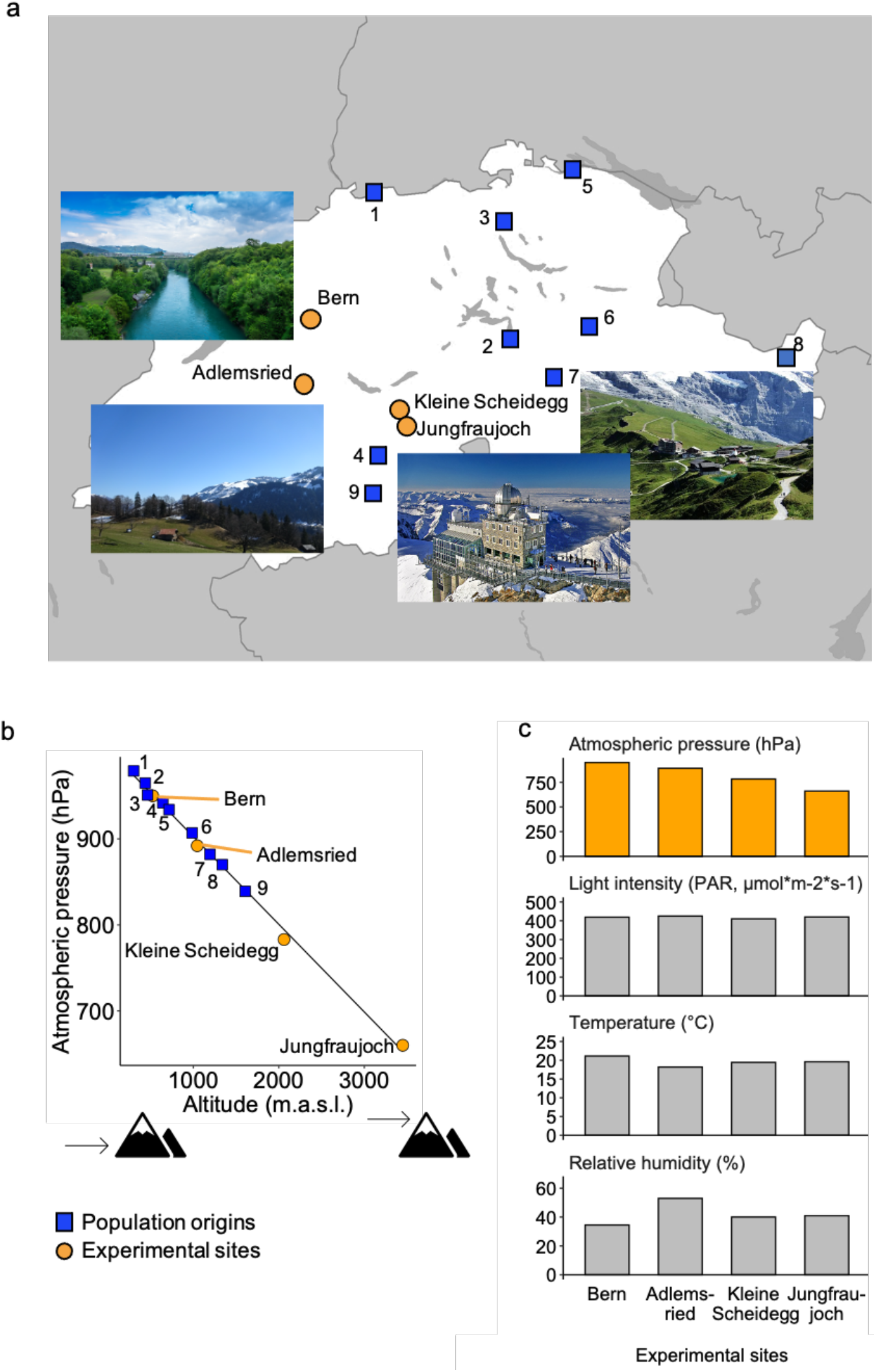
Geographical positions of the origins of the *T. officinale* populations (blue squares 1-9) and of the experimental stations (orange circles). (a) Spatial distribution of the sites across Switzerland. © Photographs: Bern, CC0 public domain; Adlemsried, Zoe Bont; Jungfraujoch, Julius Silver, CC BY-SA 4.0; Kleine Scheidegg, Grindel1, CC BY-SA 3.0. (b) Altitude and atmospheric pressure of the sites and representation of the linear relationship between these wo parameters. (c) Visualization of plant growth conditions at experimental stations, including controlled abiotic parameters (grey bars) and varying atmospheric pressure (orange bars).

### Experimental design

In order to investigate the effect of atmospheric pressure on the growth and chemical defence of *T. officinale*, we cultivated the F2 plants from the nine natural populations in four experimental stations at 526 m.a.s.l. (Bern), 1046 m.a.s.l. (Adlemsried), 2061 m.a.s.l. (Kleine Scheidegg) and 3450 m.a.s.l. (Jungfraujoch) in Switzerland (Fig. 1). Although *T. officinale* does not occur naturally in Switzerland at 3450 m.a.s.l. due to the cold climate, the plant is frequently reported at this altitude in the South American Andes (Sandoya *et al.* 2017) and is therefore not restricted in its occurrence by the associated low atmospheric pressure. In the experimental stations, the plants were grown inside under controlled light supply, temperature and relative humidity and ambient indoor air quality. This allowed us to specifically test the influence of the atmospheric pressure gradient, ranging from 950 hPa to 660 hPa (Fig. 1c).

For experiments, seeds from 6-8 mother plants (F1) per population were germinated on moist seedling substrate and transplanted into individual 1l pots filled with potting soil (5 parts field soil, 4 parts peat, 1 part sand) after 18 ± 2 days. All plants were grown under controlled conditions (25 ± 2 °C, 60 ± 5% RH, 16:8 light:dark cycle) in Bern. 24 h after transplantation, the plants were distributed among the four experimental stations, so that 6-8 plants from each of the nine populations were cultivated in each station. The temperature and relative humidity were measured hourly at all stations with RHT10-Data Logger (Extech Instruments, China). To standardize light supply, we grew the plants in rooms with reduced natural light and supplied them with LED lights (400 LED beads, LED type SMD 5730, 400 ± 20 μmol*m^2^*s^1^) placed one meter above the plants. The plants were watered three times a week and fertilized once a week.

### Assessment of plant traits and chemical analysis

After 45 days of growth in the experimental stations, we transported the plants back to Bern to analyse growth and defence traits. To assess vegetative growth, we quantified the plant biomass by measuring the dry weight of roots and shoots separately and then calculated root:shoot ratios. To study defence traits, we measured the amount of taproot latex and quantified the concentration of the latex secondary metabolites TA-G, di-PIEs and tri-PIEs.

To quantify latex traits, plants were cut 0.5 cm below the tiller and the taproot latex that was released over 20 seconds was collected and weighted. Two μl of latex was immediately transferred into 200 μl methanol for chemical analysis. The roots were carefully washed with tap water and roots and shoots were placed in a drying oven at 50°C until constant dry mass was reached. The chemical analysis of the latex metabolites was carried out as described in Bont *et al.*, 2017. Briefly, the samples were vortexed for 10 min, ultrasonicated for 10 min, centrifuged at 4 °C and 14000 rpm for 20 min and supernatants were used for further analysis. Relative concentrations of TA-G, di-PIEs and tri-PIEs were determined by injecting the latex extracts into an Acquity UPLC-PDA-MS (Waters, Milford MA, USA) with electrospray ionization in positive mode, consisting of an ultra-performance liquid chromatograph (UPLC) coupled to a photodiode array detector (PDA) and a single quadrupole mass detector (QDa). For quantification, peak areas were integrated at 245 nm for TA-G and at 275 nm for di- and tri-PIES, while concurrently recorded characteristic mass features were used to confirm compound identities (Bont *et al.* 2017). For absolute quantification of TA-G, we established an external standard curve with loganin (CAS: 18524-94-2, Sigma-Aldrich Chemie GmbH, Buchs, Switzerland) and calculated the corresponding response factor to pure TA-G. (Bont *et al.* 2017). For di-and tri-PIEs, relative concentrations were calculated separately.

### Statistical analysis

All statistical analyses were performed in R 4.0.2 (R Core Team 2017). To represent the climatic conditions associated with the different altitudes of the populations we first conducted a principal component analysis (PCA), as some of the meteorological variables of the population origins were highly correlated (Fig. S1c). This approach is widely used to analyze the impact of climate on the evolution of plant traits (Keller *et al.* 2009; Kooyers *et al.* 2015; Villaverde *et al.* 2017). We applied the function ‘prcomp’ on scaled variables to reduce dimensionality of the data and selected the axis that explained most of the cumulative variance and represented the climatic conditions associated with the different altitudes at which the populations evolved (climPCA1).

For data exploration, we calculated a correlation matrix of all parameters included in the experiment. Then, we analysed all data with linear mixed-effects models (LMEMs) (Bates 2020). The models were fit using the function ‘lmer’ from the package ‘lme4’ (Bates *et al.* 2015) with restricted maximum likelihood estimation (REML). Variables representing fixed effects were scaled and centred prior to computation to reduce nonessential multicollinearity (Iacobucci *et al.* 2016). If necessary, log-transformation was applied to the response variable to improve distribution of variance. Model assumptions were validated using ‘plotresid’ from the package ‘RVAideMemoire’ (Hervé 2018). The significances of the fixed effects were estimated using the package ‘lmerTest’ (Kuznetsova *et al.* 2017) by calculating type II analysis of variance tables with Kenward-Rodger’s approximation to degrees of freedom (Halekoh & Højsgaard 2014).

To test the overall effect of the varying atmospheric pressure of the experimental stations on plant growth and defence, we performed LMEMs separately for each plant trait (root dry weight, shoot dry weight, root:shoot ratio, taproot latex, TA-G, di-PIEs, tri-PIEs). We used the mean value per trait per population per station as response variable, tested the fixed effect of the atmospheric pressure of the experimental station (‘P_Station_’) and included ‘(1|Population)’ as random effect and grouping factor to allow for varying intercepts between populations. We performed a similar analysis to assess the overall effect of the climatic conditions associated with the different altitudes of the population origins on plant growth and defence by testing the fixed effect of ‘climPCA1’ and including ‘(1|Station)’ as random effect and grouping factor to allow for varying intercepts between experimental stations.

In order to further investigate interacting effects of the climatic conditions associated with different altitudes of the population origins (genetic effects) and the atmospheric pressure of the experimental stations (environmental effects) on the plants, we next conducted a full model analysis for each plant trait, including the abiotic environment of the stations as covariate. Since temperature and relative humidity were highly correlated (Pearson’s *r* = −0.95, Fig. S3), we used only temperature as covariate in the model. Trait values of individual plants were used as response variables, with ‘(1|Population)’ as random effect to allow the calculation of separate intercepts for each population. Concurrently, the impact of ploidy was tested by adding the ploidy level (diploid or triploid) of each plant as fixed effect. The full model syntax was the following: plant trait ~ climPCA1 × P_Station_ + P_Station_ × temperature + P_Station_ × Ploidy + (1|Population). As expected, temperature influenced all experimental parameters (LMEMs: Temp, *P* < 0.05, Table 1), but did not interact with atmospheric pressure (LMEMs: P_Station_ × Temp, *P* < 0.05, Table 1). A confounding effect of residual variation in temperature on the detected effects of atmospheric pressure is, therefore, unlikely. To visualize the significant interactive effect of ‘climPCA1 × P_Station_’ on TA-G and on di-PIEs concentration, we then used the package ‘effects’ (Fox *et al.* 2019) for model prediction with unscaled and uncentered fixed effects, excluding the non-significant interaction of ‘P_Station_ × temperature’ from the model. Likewise, we predicted and visualized the nonsignificant effects of P_Station_ × Ploidy on all measured plant traits. All results were visualized using ‘ggplot2’ (Wickham 2016) and ‘effects’ (Fox *et al.* 2019).

**Table 1:** Effects of experimental atmospheric pressure (P_Station_), of climatic conditions associated with different altitudes of population origins (climPCA1), of the plant’s ploidy level (Ploidy) and of selected interactions on performance and latex profile of *T. officinale* are shown. Temperature of experimental station (Temp) is included as control variable. Results of full mixed-effects model analyses are displayed separately for each performance parameter (root dry weight, shoot dry weight, root:shoot ratio, amount of latex) and for each class of latex secondary metabolites. Significances of fixed effects were assessed by *F*-tests. Estimated *F*-values_(NumDF, DenDF)_ are shown. Levels of statistical significance are indicated with asterisks (\*\*\**P* <0.001; \*\**P* <0.01; \**P* <0.05; (*) *P* < 0.1). TA-G: taraxinic acid ß-D-glucopyranosyl ester; di-PIEs: di-4-hydroxyphenylacetate inositol esters; tri-PIEs: tri-4-hydroxyphenylacetate inositol esters.

|  | Root | Shoot | Root:shoot | Latex | TA-G | di-PIEs | tri-PIEs<sup>1</sup> |
| --- | --- | --- | --- | --- | --- | --- | --- |
| climPCA1 | 0.83<sub>(1,6)</sub> | 2.08<sub>(1,6)</sub> | 0.00<sub>(1,6)</sub> | 5.20<sub>(1,6)</sub><br>(*) | 11.00<sub>(1,6)</sub><br>* | 2.05<sub>(1,6)</sub> | 0.35<sub>(1,6)</sub> |
| $P_{\text{Station}}$ | 10.75<sub>(1,209)</sub><br>** | 1.29<sub>(1,209)</sub> | 18.22<sub>(1,209)</sub><br>*** | 0.01<sub>(1,209)</sub> | 14.12<sub>(1,211)</sub><br>*** | 0.01<sub>(1,211)</sub> | 5.60<sub>(1,211)</sub><br>* |
| Temp | 6.26<sub>(1,210)</sub><br>* | 4.16<sub>(1,210)</sub><br>* | 1.62<sub>(1,209)</sub> | 11.41<sub>(1,210)</sub><br>*** | 4.17<sub>(1,211)</sub><br>* | 8.03<sub>(1,211)</sub><br>** | 8.21<sub>(1,211)</sub><br>** |
| Ploidy | 0.06<sub>(1,108)</sub> | 0.92<sub>(1,107)</sub> | 0.38<sub>(1,143)</sub> | 0.06<sub>(1,114)</sub> | 0.23<sub>(1,101)</sub> | 1.36<sub>(1,95)</sub> | 0.38<sub>(1,209)</sub> |
| climPCA1<br>x $P_{\text{Station}}$ | 0.72<sub>(1,209)</sub> | 2.47<sub>(1,209)</sub> | 0.40<sub>(1,209)</sub> | 0.08<sub>(1,209)</sub> | 7.26<sub>(1,211)</sub><br>** | 6.49<sub>(1,211)</sub><br>* | 2.06<sub>(1,211)</sub> |
| $P_{\text{Station}}$ x<br>Temp | 0.49<sub>(1,210)</sub> | 0.13<sub>(1,210)</sub> | 1.33<sub>(1,210)</sub> | 3.32<sub>(1,210)</sub><br>(*) | 0.27<sub>(1,212)</sub> | 0.00<sub>(1,212)</sub> | 0.09<sub>(1,211)</sub> |
| $P_{\text{Station}}$ x<br>Ploidy | 0.67<sub>(1,209)</sub> | 1.75<sub>(1,209)</sub> | 1.08<sub>(1,209)</sub> | 0.50<sub>(1,209)</sub> | 0.91<sub>(1,211)</sub> | 0.48<sub>(1,211)</sub> | 3.30<sub>(1,211)</sub> |

<sup>1</sup> log-transformed

## Results

### Validation of experimental setup

We first tested whether our efforts to standardize environmental parameters across experimental sites were successful. We found that indoor temperature and relative humidity varied slightly between sites (Fig. 1c), but neither variable was correlated to atmospheric pressure (Fig. S3). As plants were also grown using identical potting soil, watering, lighting and comparable indoor environments with respect to air pollution, we estimate that major abiotic parameters were successfully standardized to allow for an assessment of the role of atmospheric pressure on *T. officinale* (see discussion for a critical assessment of this aspect). By growing *T. officinale* plants from seeds of diploid and triploid populations originating from different altitudes, we tested (i) the effect of atmospheric pressure on growth and defence; (ii) the impact of population origin on plant responses to atmospheric pressure (as an indicator for local adaptation to atmospheric pressure); and (iii) the importance of a plant’s cytotype (diploid vs. triploid) on its capacity to grow under different atmospheric pressure, as a test of the hypothesis that low atmospheric pressure restricts the establishment of triploids at high altitudes. To this end, we first describe environmental (Fig. 2) and heritable contributions (Fig. 3) to variation in defence and growth independently. Using a full model, we then address adaptive phenotypic plasticity (Table 1, Fig. 4) and differences between cytotypes (Table 1, Fig. 5).

**Figure 2:**
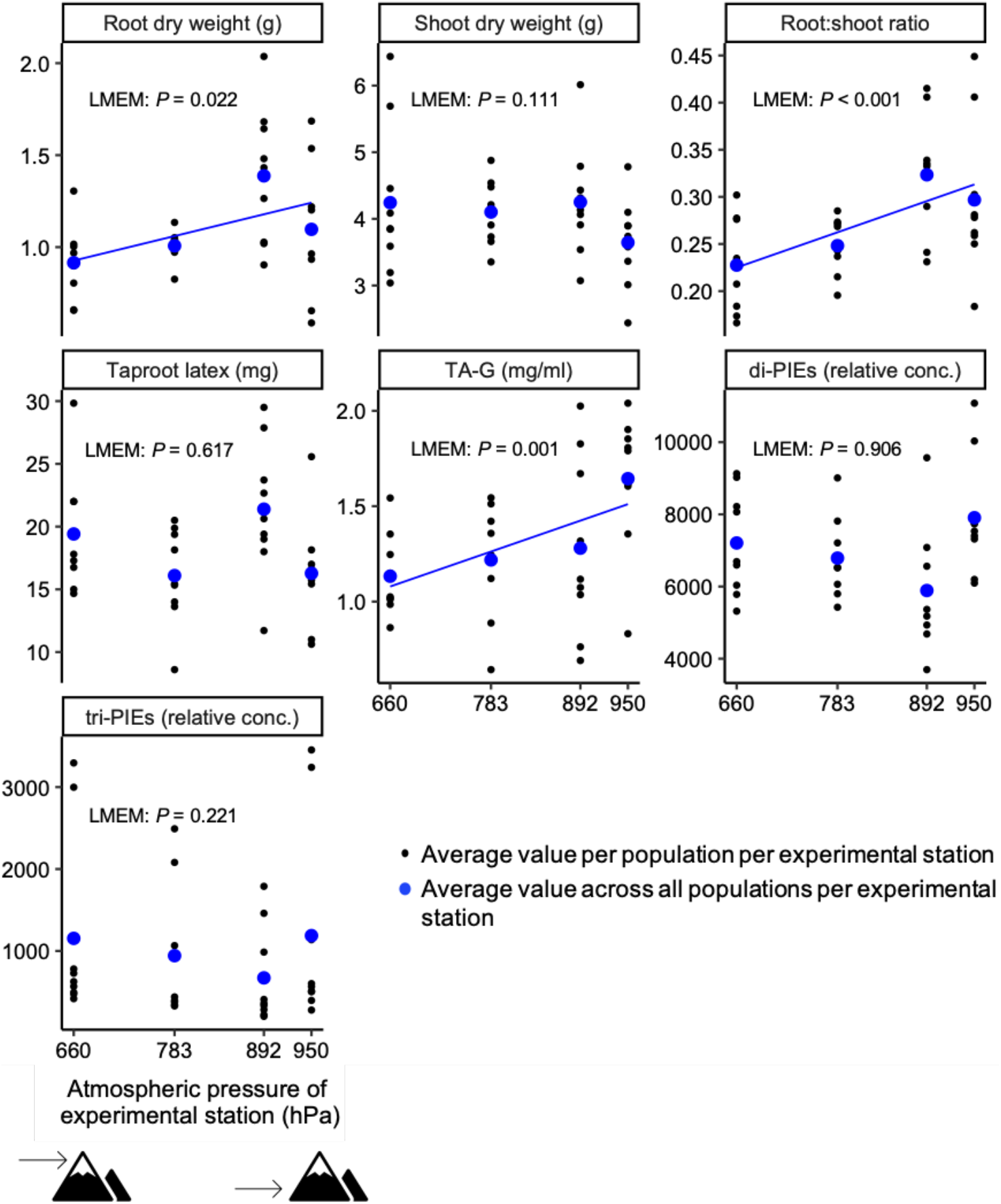
Overall effect of atmospheric pressure of experimental station on mean plant performance parameters (root dry weight, shoot dry weight, root:shoot ratio, taproot latex) and chemical composition of latex (TA-G, di-PIEs, tri-PIEs). Blue dots represent average values across all populations per experimental station, while black dots represent average values per population per experimental station (*N* = 5-8 per population and station). The significance of the effect was tested with linear mixed-effects models (LMEM) and corresponding *P*-values are displayed. Blue lines indicate statistically significant effects (*P* < 0.05). TA-G: taraxinic acid ß-D-glucopyranosyl ester; di-PIEs: di-4-hydroxyphenylacetate inositol esters; tri-PIEs: tri-4-hydroxyphenylacetate inositol esters.

**Figure 3:**
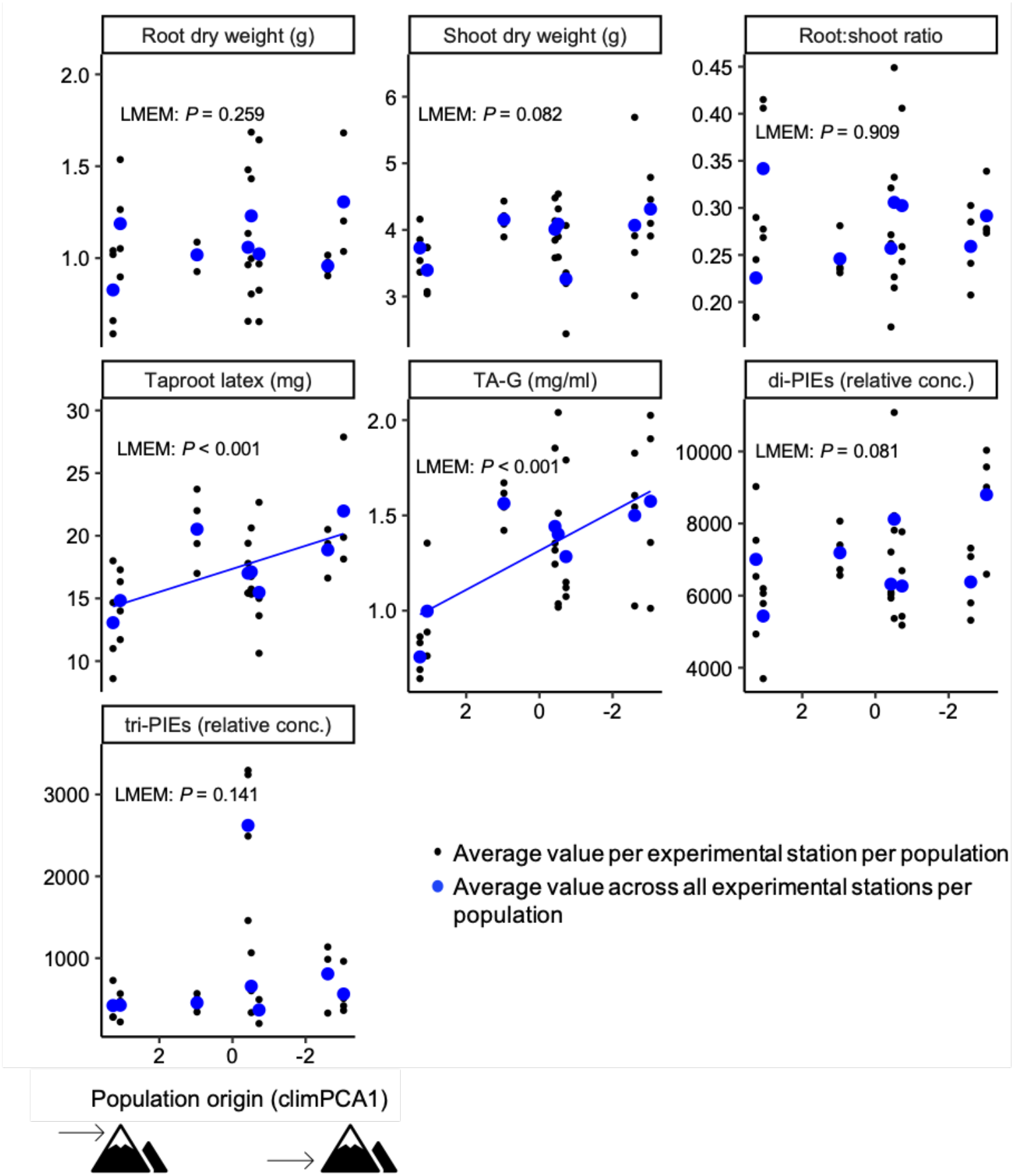
Overall effect of climatic conditions associated with different altitudes of population origins (climPCA1) on mean plant performance parameters (root dry weight, shoot dry weight, root:shoot ratio, taproot latex) and chemical composition of latex (TA-G, di-PIEs, tri-PIEs). climPCA1 represents a climatic gradient ranging from high altitude environment to low altitude environment, which is illustrated by icons. Blue dots represent average values per population across all experimental stations, while black dots represent average values per population per experimental station (*N* = 5-8 per population and station). The significance of the effect was tested with linear mixed-effects models (LMEM) and corresponding *P*-values are displayed. Blue lines indicate statistically significant effects (*P* < 0.05). TA-G: taraxinic acid ß-D-glucopyranosyl ester; di-PIEs: di-4-hydroxyphenylacetate inositol esters; tri-PIEs: tri-4-hydroxyphenylacetate inositol esters.

**Figure 4:**
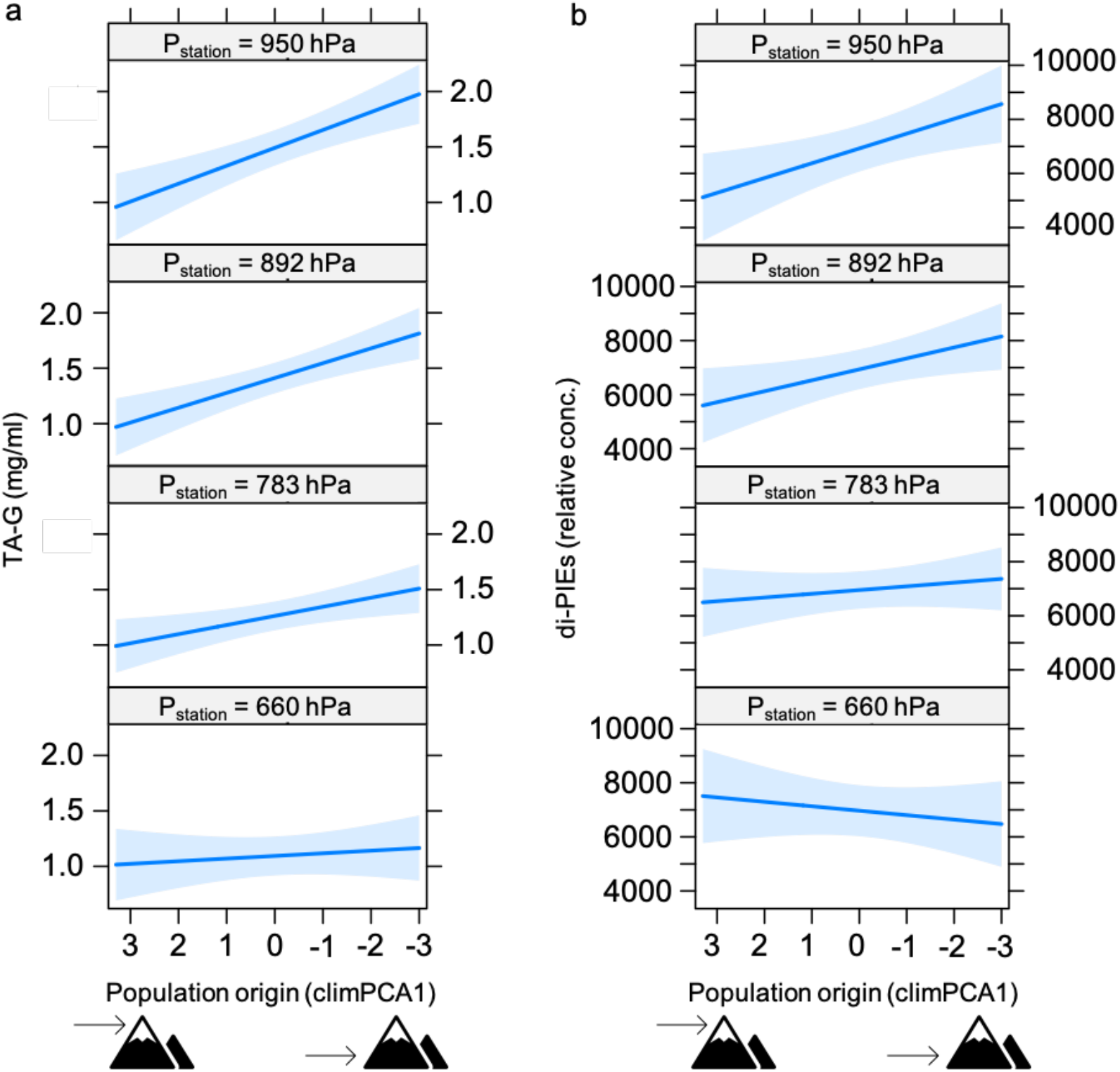
Visualization of the significant interaction effect of the climatic conditions associated with different altitudes of the population origins (climPCA1) on TA-G concentration (a) and on concentration of di-PIEs (b) in the latex depending on the atmospheric pressure of the experimental station. Blue lines indicate predicted slopes from the mixed-effects model, with 95% confidence interval shaded. TA-G: taraxinic acid ß-D-glucopyranosyl ester; di-PIEs: di-4-hydroxyphenylacetate inositol ester.

**Figure 5:**
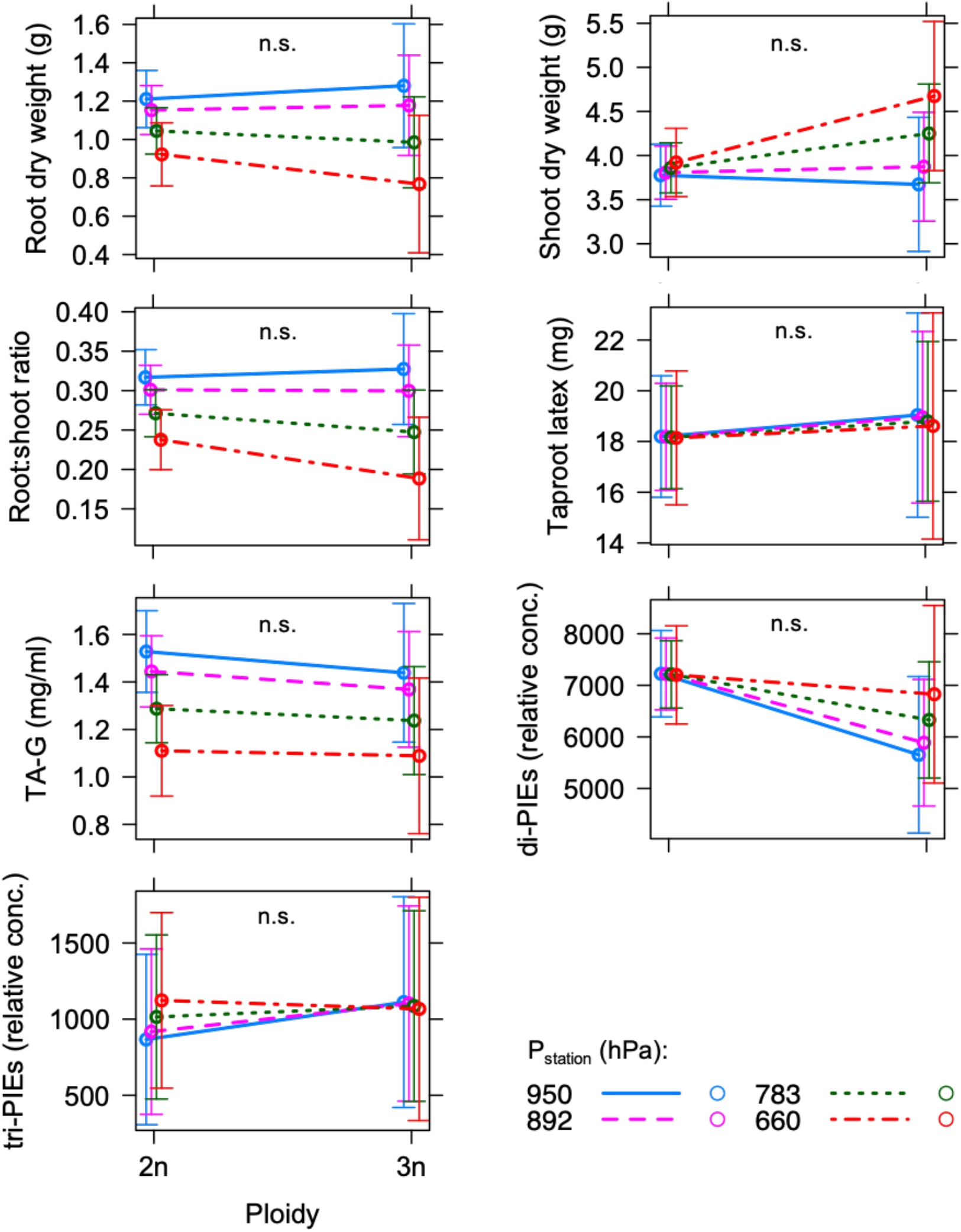
Visualization of influence of ploidy on plant parameters depending on atmospheric pressure of experimental station. Predicted mean values with standard errors from mixed-effects models are shown. None of the plant parameters differ statistically significantly between diploid and triploid plants (n.s.: *P* > 0.05). 2n: diploid; 3n: triploid; TA-G: taraxinic acid ß-D-glucopyranosyl ester; di-PIEs: di-4-hydroxyphenylacetate inositol esters; tri-PIEs: tri-4-hydroxyphenylacetate inositol esters.

### Atmospheric pressure modulates root growth and defences

Across all populations and irrespective of population origin and cytotype, *T. officinale* plants showed distinct phenotypes as a function of atmospheric pressure at the different experimental sites (Fig. 2). Both root growth and root:shoot ratio were overall lower at higher altitudes with lower atmospheric pressure (LMEM, *P* = 0.022, Fig. 2 resp. LMEM, *P* < 0.001, Fig. 2). By contrast, no significant effects on shoot growth and on the amount of exuded taproot latex were observed (LMEMs, *P* > 0.05, Fig. 2). The concentration of the defence metabolite TA-G in the taproot latex was reduced in plants growing under lower atmospheric pressure (LMEM, *P* = 0.001, Fig. 2). The concentrations of di-PIEs and tri-PIEs did not vary significantly with atmospheric pressure (LMEMs, *P* > 0.05, Fig. 2). Thus, growth under low atmospheric pressure is specifically associated with lower root growth and lower accumulation of a defensive sesquiterpene lactone in *T. officinale*.

### Climatic history associated with altitude shapes heritable variation in root defences

To characterize the climatic conditions under which the *T. officinale* populations evolved, we conducted a principal component analysis (PCA) and then scaled variables to reduce the dimensionality of the data. The first two axes (climPCA1 and climPCA2) together explained 90% of the cumulative variance in climate parameters (Fig. S1a). 54.3% of the variance was explained by the first axis (climPCA1) which mainly represents variation in temperature-related parameters and atmospheric pressure (Fig. S1a, Fig. S1c). ClimPCA1, but not climPCA2 was highly correlated with the altitude of the population origin (Pearson’s *r* = 0.941, *P* < 0.001, Fig. S1b). We therefore used climPCA1 to represent the climatic conditions associated with the different altitudes of the populations.

Across experimental stations, we found no significant association between climPCA1 and *T. officinale* growth (LMEMs, *P* > 0.05, Fig. 3). However, climPCA1 was correlated to the amount of taproot latex and the concentration of TA-G, with populations originating from lower altitudes producing more latex and more TA-G than populations collected at high altitudes (LMEMs, *P* < 0.001, Fig. 3). No clear effects were observed for di- and tri-PIEs (LMEMs, *P* > 0.05, Fig. 3). Thus, climatic conditions that are correlated with altitude are associated with heritable differences in the production and chemical composition of root latex, with plant originating from higher altitudes producing less latex and defensive sesquiterpene lactones.

### Root defences show patterns of adaptation to high atmospheric pressure

In a next step, we constructed a full model to detect interactions between the climatic history of the populations (climPCA1) and the atmospheric pressure of the experimental sites at which the plants were grown (P_Station_). The full model confirmed that the concentration of TA-G in the latex is influenced by climPCA1 (LMEM: climPCA1, *P* = 0.017, Table 1, Fig. 3), and that atmospheric pressure at the experimental sites is associated with changes in root growth (LMEM: P_Station_, *P* = 0.001, Table 1, Fig. 2), root:shoot ratio (LMEM: P_Station_, *P* < 0.001, Table 1, Fig. 2) and TA-G concentration in root latex (LMEM: P_Station_, *P* < 0.001, Table 1, Fig. 2). In addition, we detected a negative effect of atmospheric pressure on the concentration of tri-PIEs (LMEM: P_Station_, *P* = 0.019, Table 1, Fig. 2). We detected no significant interactions between the climatic history of the populations (climPCA1) and atmospheric pressure at the experimental sites for root or shoot growth (LMEM: climPCA1 × P_Station_, *P* = 0.05, Table 1). However, a significant interaction was observed for TA-G concentration (LMEM: climPCA1 × P_Station_, *P* = 0.008, Table 1), indicating natural selection for phenotypic variability of TA-G production under different atmospheric pressure. Closer inspection of the data revealed that TA-G concentrations in populations originating from low altitude environments were more strongly influenced by atmospheric pressure than in populations from high altitude environments (Fig. 4a). Under high atmospheric pressure, populations from low altitude environments produced significantly more TA-G than populations from high altitude environments (Fig. 4a). With decreasing atmospheric pressure, this difference disappeared, leading to similar TA-G concentrations at 660 hPa. A similar pattern, albeit with more variability, was observed for di-PIEs (LMEM: climPCA1 × P_Station_, *P* = 0.012, Table 1, Fig. 4b). These results are suggestive of adaptation of root defence expression of low altitude populations to high atmospheric pressure.

### The performance of triploid plants is not constrained by resistance to atmospheric pressure

To test whether plant cytotype influences the capacity of *T. officinale* to grow at different atmospheric pressures, we tested for interactions between cytotype and atmospheric pressure at the different experimental sites. Across experimental sites, we did not detect any significant differences between diploid and triploid plants for any of the measured parameters (LMEMs: Ploidy, *P* > 0.05, Table 1, Fig. 5). We also found no significant interactions between cytotype and atmospheric pressure (LMEMs: P_Station_ × Ploidy, *P* > 0.05, Table 1, Fig. 5). Thus, the hypothesis that low atmospheric pressure restricts the establishment of triploids at high altitudes is not supported by our data.

## Discussion

The sum of environmental factors varying with changing altitude can simultaneously impose multiple selection pressure on plant traits, making it difficult to disentangle single effects. By growing several *T. officinale* populations in controlled environments, our study provides evidence that atmospheric pressure influences plant root growth and chemical defence irrespective of the cytotype level of the plants and shows that the natural habitats of the populations shape the potential for phenotypic variability in response to varying atmospheric pressure. Here, we discuss our findings in an eco-evolutionary and plant-physiological context.

Although many plants seem to be capable to grow vegetatively at pressures even below 25 kPa (Richards *et al.* 2006), they suffer from stress associated with hypoxia and desiccation and in turn develop responses and adaptations, which are often organ-specific (Zhou *et al.* 2017). In environments with reduced O_2_ and CO_2_ availability, roots may react more strongly to oxygen deficiency because, as heterotrophic organs, they are highly dependent on oxygen for mitochondrial energy production, while shoots, as autotrophic plant organs, may be restricted in photosynthesis due to the reduced CO_2_ diffusion rate (Mustroph *et al.* 2014). Our results show that root but not shoot growth of *T. officinale* is reduced in low atmospheric pressure environments, resulting in a decrease of the root:shoot ratio. In our experimental stations, the reduction in atmospheric pressure is associated with naturally decreased levels of atmospheric gases including a reduced partial pressure of O_2_, and we assume that the resulting mild hypoxia restricts root growth of *T. officinale*. This finding is in line with previous studies showing that roots are particularly sensitive to reduced O_2_ partial pressure (He *et al.* 2007; Tang *et al.* 2015), whereas several studies with plants cultivated at reduced atmospheric pressure, but with a partial pressure of O_2_ experimentally maintained at the same level as at ambient pressure, found no reduction in root biomass (Iwabuchi *et al.* 1996; Spanarkel & Drew 2002; Levine *et al.* 2008).

Exposure to low atmospheric pressure also affects plant chemistry (He *et al.* 2013; Levine *et al.* 2008; Zhou *et al.* 2017). In our study we detected a decline in the concentration of the secondary metabolite TA-G in the latex of *T. officinale* with decreasing atmospheric pressure. TA-G is constitutively produced by the plant, acts repellent against root feeders and therefore defends the plant against herbivores (Huber *et al.* 2016b; Bont *et al.* 2017). To reduce the costs associated with constitutively produced chemical defenses (Neilson *et al.* 2013), plants may use abiotic conditions as external stimuli to adjust the level of defensive secondary metabolites to the expected herbivore pressure. Temperature, for instance, is assumed to be a good indicator for herbivore attack in the field, because herbivore appearance and activity is often modulated by temperature (Bale *et al.* 2002). In our previous work, we found evidence that *T. officinale* can use seasonal temperature variation to synchronize deployment of chemical defenses with expected herbivore attack intensity in the field, indicating an important role of abiotic conditions in fine-tuning the level of constitutively produced defensive metabolites (Huang *et al.* 2019). With increasing altitude, it is often assumed that herbivore pressure decreases (Moreira *et al.* 2018). *T. officinale* might thus use low atmospheric pressure as an indicator of high-altitude growth conditions and associated expected lower herbivore pressure, and the observed decrease in TA-G production might be a fine-tuning of defense deployment to reduce costs and maximize plant fitness. However, these are highly speculative conclusions and require further investigations.

Our previous work has not only shown that TA-G is involved in herbivore defense in *T. officinale*, but it has also shown an association of TA-G with the climatic history of the plants’ natural habitats, suggesting an additional role of TA-G in abiotic stress management (Bont *et al.* 2020). The climatic parameters associated with TA-G production included sun and rain intensity and were related to the latitude but not to the altitude of the natural populations – plants from the rainier North of Switzerland produced more TA-G than plants from the sun-intense regions in the South of Switzerland (Bont *et al.* 2020), leading to the hypothesis that TA-G may passively or actively be involved in moisture regulation. At low atmospheric pressure, water evaporation is increased, and evidence exists that when plants are grown under such conditions, their perceptual mechanisms of water movement are altered even though the plants are fully hydrated and do not experience actual desiccation (Paul *et al.* 2004; Zhou *et al.* 2017). The decline in TA-G concentration with decreasing atmospheric pressure observed in our study could therefore be a direct or indirect consequence of increased evapotranspiration and would then support an involvement of TA-G in moisture regulation.

The capacity of phenotypic plasticity allows a plant with a given genotype to adjust its phenotype to the demands of contrasting environments (Nicotra *et al.* 2010; Sultan 2000). In our experiments, the employed atmospheric pressure gradient was associated with measurable phenotypic plasticity, which again emphasizes the importance of atmospheric pressure for plant growth and development (Paul & Ferl 2006). For the latex metabolites TA-G and di-PIEs, plastic responses differed among populations, indicating heritable within-species variation for plasticity in these traits, likely shaped by the climatic histories of the populations. In high atmospheric pressure environments, plants from low altitude environments produced more of the metabolites than plants from high altitude environments, whereas in low atmospheric pressure environments, the difference between the populations vanished. The capacity of low altitude plants to increase production of TA-G and di-PIEs when grown under low altitude atmospheric pressure could benefit these plants and may be an adaptive trait, as latex metabolites have defensive functions against root feeders (Huber *et al.* 2016a, b; Bont *et al.* 2017) and herbivore pressure is often expected to increase with decreasing altitude (Rasmann *et al.* 2014; Moreira *et al.* 2018). These findings are consistent with previous studies showing that within species, constitutive defenses often decrease with increasing altitude (Bakhtiari *et al.* 2019; Buckley *et al*. 2019; Meyer & Carlson 2001; Pellissier *et al.* 2014). Our results further suggest that climatic conditions characterizing low altitude environments select for genotypes with high plasticity in latex metabolite production, although whether the observed plasticity is a passive or active plastic response (van Kleunen & Fischer 2005) and whether its expression increases plant fitness remain to be elucidated.

Along altitudinal transects, diploid sexual *T. officinale* predominate over triploid asexual individuals at higher altitudes (Calame & Felber 2000), while along latitudinal gradients, triploids predominate over diploids at higher latitudes (Van Dijk *et al.* 2003). In our study, we tested the hypothesis that the establishment of triploids at high altitudes is restricted by low atmospheric pressure. However, we found no evidence for a disadvantage of triploids at low atmospheric pressure, as diploids and triploids did not differ in any of the measured plant traits at different atmospheric pressures. We conclude that atmospheric pressure is unlikely to influence geographic parthenogenesis of *T. officinale* and that other factors, for example recolonization patters after glaciation (Calame & Felber 2000; Kearney 2005), might have influenced the spatial distribution of diploids and triploids along certain transects.

Studies comparing different species have shown that plants growing in high altitude environments have developed adaptations to cope with the challenges of these environments (Halbritter *et al.* 2018) and that some of these adaptations, for example changes in stomatal development, are directly related to atmospheric pressure (Kammer *et al.* 2015). Our study confirms that atmospheric pressure is an important abiotic factor that influences plastic responses in plant growth and development and shows that across *T. officinale* populations, exposure to varying atmospheric pressures evokes heritable responses in growth and defense traits - responses that are shaped by the climatic conditions of the natural habitats of the populations. Given the non-negligible impact of atmospheric pressure on the expression and likely also on the evolution of plant traits, we suggest that atmospheric pressure should be included by default as an abiotic factor when studying plant variation along altitudinal gradients to prevent results from being obscured by possibly unrecognized effects of atmospheric pressure variation. This may be particularly important when altitudinal gradients are used to surrogate climate change (Carlyle *et al.* 2014; Frei *et al.* 2014; Michalet *et al.* 2014), since atmospheric pressure, unlike temperature or precipitation, is not expected to change under global warming.

An underlying assumption of our work is that we successfully standardized or randomized environmental parameters other than atmospheric pressure across the different experimental stations, thus allowing us to infer effects of this factor. Indeed, our experiment allowed us to control and/or randomize all major abiotic and biotic environmental parameters, including soil structure and composition, water supply, humidity, temperature as well as light quality and quantity and air pollution. Pests and pathogens were not observed on our plants. Another factor that depends on altitude and co-varies with atmospheric pressure is gravity. Gravity decreases with altitude, resulting in a delta of 0.01 m/s^2^ between Bern and the Jungfraujoch. Whether such a small change in gravity has any measurable impact on plants is unknown. Plant gravitropism depends on sensing inclination rather than gravitational force (Chauvet *et al.* 2016), and effects of reduced gravitational force are typically only observed below 0.3 *g* (Kiss *et al.* 2019). Thus, we infer that the effects observed in our study are the result of changes in atmospheric pressure rather than other environmental factors. Further experiments with pressure chambers (Paul *et al.* 2004) could be used as a future approach to confirm the patterns observed in our study.

Our results emphasize that within species, plants from different populations respond differently to varying atmospheric pressure, especially in the production of secondary metabolites. For successful cultivation of plants in extraterrestrial habitats as food source, it therefore may be worthwhile to screen various populations of the species of interest under low atmospheric pressure to detect resistant populations with secondary metabolite profiles suitable for human nutrition.

## Acknowledgements

We thank Marc Pfander for his support with statistical analysis. We are grateful to Andrea Bonini for his help with field and lab work and to the gardeners of the University of Bern for taking care of the plants. We thank Hildegard Hinni, Annerös Erb, André Hofer, Christoph Steiner, Joan Fischer and Martin Fischer for watering the plants during the experiments at the research stations in Adlemsried, Kleine Scheidegg and Jungfraujoch. We thank the Kleine Scheidegg station of the Swiss Federal Railways to provide the room to perform the experiment. We thank the International Foundation High Altitude Research Station Jungfraujoch and Gornergrat (HFSJG) for the opportunity to conduct experiments. Meteorological data was provided by MeteoSwiss, the Swiss Federal Office for Meteorology and Climatology. This study was supported by the Swiss National Science Foundation (Grant No. 153517) and the Seventh Framework Programme for Research and Technological Development of the European Union (FP7 MC-CIG 629134).

## Notes

### Competing Interest Statement

The authors have declared no competing interest.

