## Supplementary material for "Adaptations and responses of the common dandelion to low atmospheric pressure in high altitude environments"

### Supporting information

**Table S1** Descriptive parameters of the nine natural *T. officinale* populations included in the experiment. Location of the populations (latitude, longitude, altitude), abbreviation of the corresponding weather station (WS), altitude of the weather station, long-term average of atmospheric pressure measured by the weather station and number of diploid and of triploid mother plants included in the experiment are shown.

| Popul ation | Latitude (°N) | Longitude (°E) | Altitude (m.a.s.l.) | Weather station (WS) | WS: Altitude (m.a.s.l.) | WS: atmos. pressure (hPa) | Mother plants: diploid | Mother plants: triploid |
|---|---|---|---|---|---|---|---|---|
| 1 | 47°32′39″ | 7°47′54″ | 302 | RHF | 300 | 973 | 1 | 6 |
| 2 | 46°53′25″ | 8°37′12″ | 437 | ALT | 438 | 965 | 7 | 0 |
| 3 | 47°23′24″ | 8°35′55″ | 465 | SMA | 556 | 951 | 6 | 2 |
| 4 | 46°17′58″ | 7°51′31″ | 643 | VIS | 639 | 942 | 7 | 0 |
| 5 | 47°39'06″ | 9°01'53″ | 716 | HAI | 718 | 934 | 3 | 3 |
| 6 | 46°55′02″ | 9°10′15″ | 986 | ELM | 958 | 907 | 5 | 0 |
| 7 | 46°42′24″ | 8°51′13″ | 1196 | DIS | 1197 | 882 | 8 | 0 |
| 8 | 46°47′50″ | 10°17′14″ | 1339 | SCU | 1303 | 870 | 6 | 1 |
| 9 | 46°11′43″ | 7°50′13″ | 1607 | GRC | 1605 | 839 | 6 | 0 |

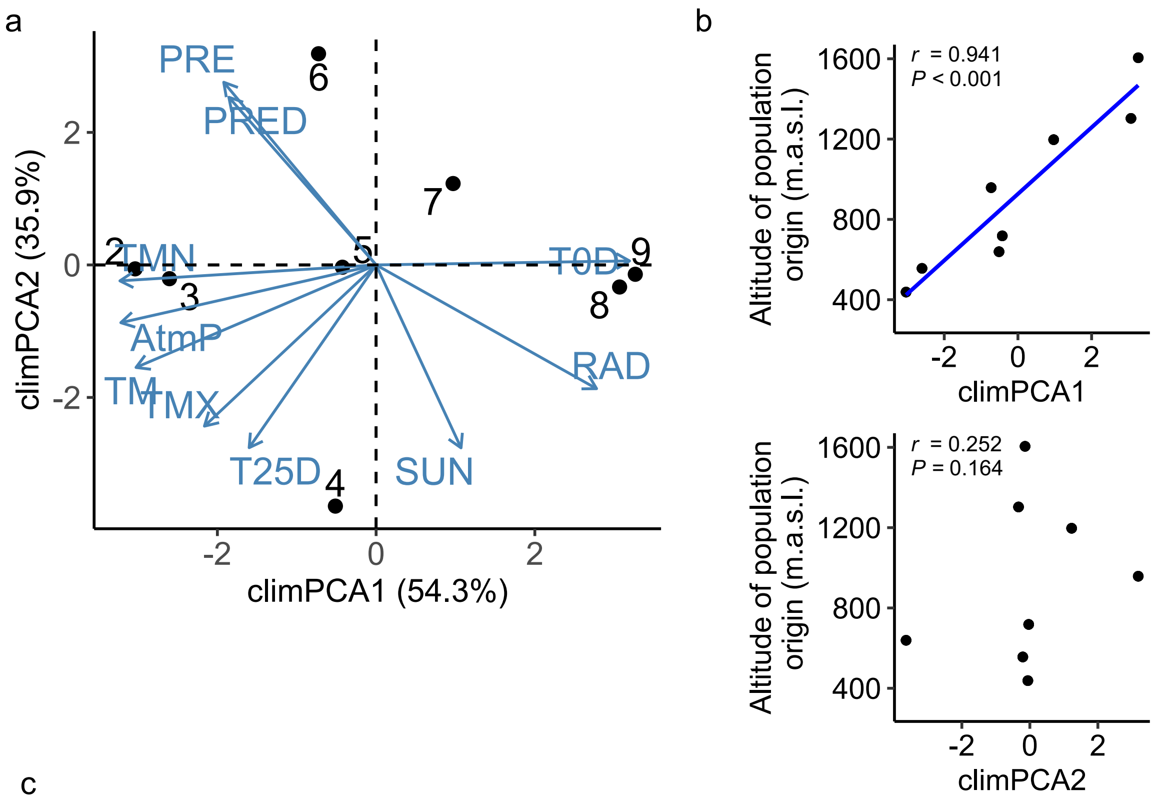


c

| Climate variable | Description | climPCA1 | climPCA2 |
|---|---|---|---|
| AtmP | Annual mean of atmospheric pressure | -0.404 | -0.135 |
| PRE | Annual precipitation | -0.241 | 0.426 |
| PRED | Days per year with precipitation | -0.233 | 0.391 |
| RAD | Annual mean of global radiation | 0.349 | -0.289 |
| SUN | Annual sunshine duration | 0.134 | -0.427 |
| TM | Annual mean temperature | -0.380 | -0.239 |
| TMN | Annual mean of the minimal temperature per day | -0.406 | -0.037 |
| TMX | Annual mean of the maximal temperature per day | -0.272 | -0.376 |
| T0D | Days per year with minimal temperature below 0°C | 0.401 | 0.009 |
| T25D | Days per year with maximal temperature over 25°C | -0.201 | -0.426 |

**Figure S1** Principal component analysis of the climatic conditions of the population origins. (a) Biplot of the analysis showing the first two axes (climPCA1, climPCA2), which explain 90% of cumulative variance. Blue arrows represent climatic conditions and black dots represent populations. (b) Pearson correlations of climPCA1 and climPCA2 with altitude of population origin. Populations are represented with black dots. Significant linear correlation line and *r*- and *P*-values of correlations are shown. (c) Description of variables used in the analysis, representing climatic conditions of the population origins. Correlations of climate variables with the first two principal comonents (climPCA1, climPCA2) of the analysis are shown.

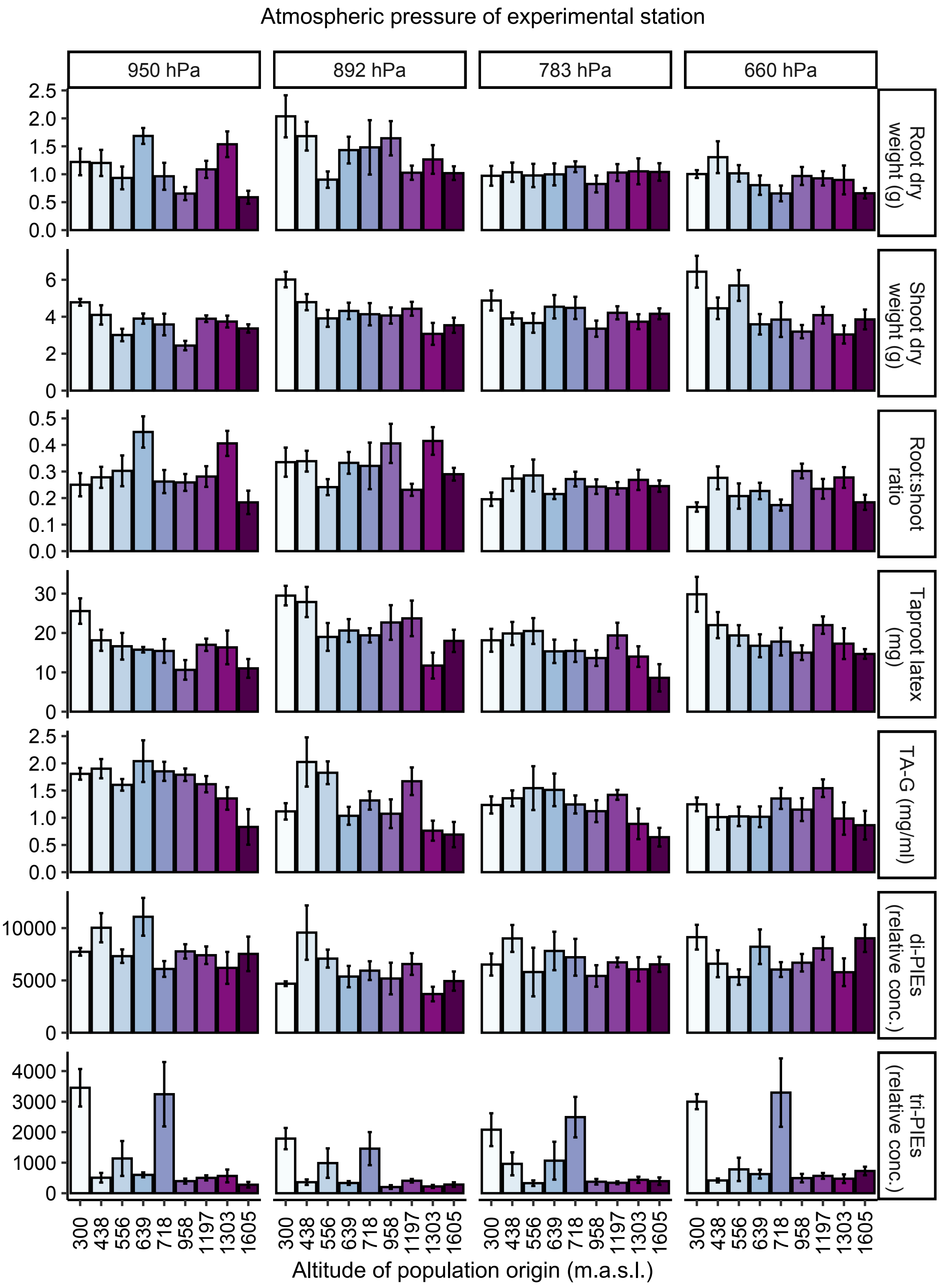


**Figure S2:** Plant performance parameters (root dry weight, shoot dry weight, root:shoot ratio, taproot latex) and concentrations of root latex secondary metabolites (TA-G, di-PIEs, tri-PIEs) are displayed separately for each *T. officinale* population growing under the atmospheric pressure of each experimental station. Bars represent population means ($N$ = 5-8) and standard errors. Bars with the same color correspond to values from the same population. TA-G: taraxinic acid ß-D-glucopyranosyl ester; di-PIEs: di-4-hydroxyphenylacetate inositol esters; tri-PIEs: tri-4-hydroxyphenylacetate inositol esters.

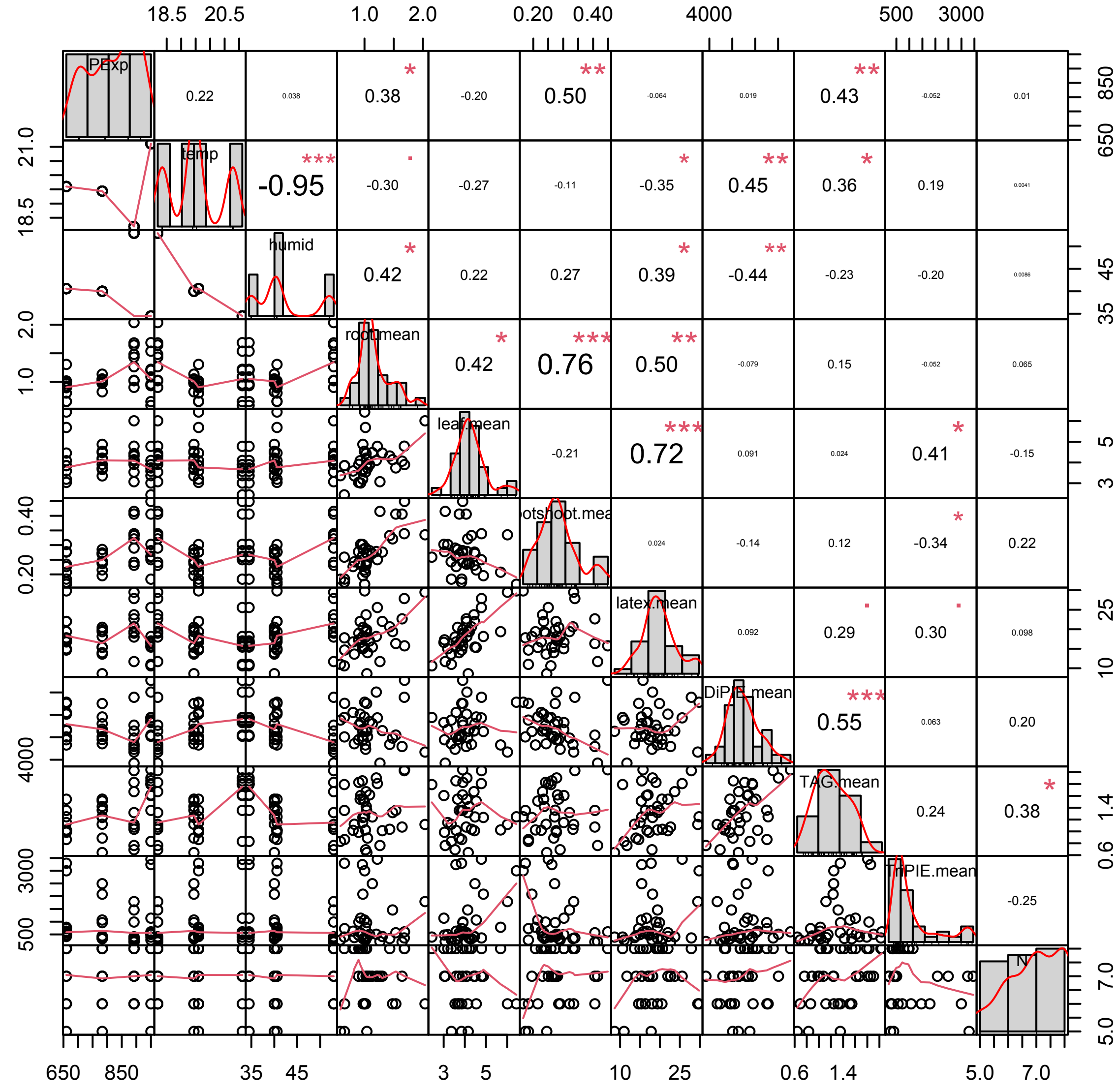


**Figure S3:** Correlation matrix of measured variables and environmental factors. PExp: Atmospheric pressure of experimental station; temp: Mean temperature of experimental station; humid: mean relative humidity of experimental station; root.mean: root dry weight; leaf.mean: shoot dry weight; root:shoot.mean: root:shoot ratio; latex.mean: taproot latex; DiPIE.mean: concentration of di-PIEs (di-4-hydroxyphenylacetate inositol esters) in the latex, TAG.mean: concentration of TAG (taraxinic acid ß-D-glucopyranosyl ester) in the latex; TriPIE.mean: concentration of tri-PIEs (tri-4-hydroxyphenylacetate inositol esters) in the latex; PC1: climatic conditions associated with different altitudes of population origins.
